# Cross-Platform Omics Prediction procedure: a game changer for implementing precision medicine in patients with stage-III melanoma

**DOI:** 10.1101/2020.12.09.415927

**Authors:** Kevin Y.X. Wang, Gulietta M. Pupo, Varsha Tembe, Ellis Patrick, Dario Strbenac, Sarah-Jane Schramm, John F. Thompson, Richard A. Scolyer, Samuel Mueller, Garth Tarr, Graham J. Mann, Jean Y.H. Yang

## Abstract

There is no consensus methodology that can account for the variation in omics signatures when they are acquired across different platforms and times. This poses a significant barrier to the implementation of valuable biomarkers into clinical practice. We present a novel procedure (Cross-Platform Omics Prediction) that accounts for these variations and demonstrate its utility in three risk models for different diseases that is suitable for prospective and multi-centre clinical implementation.

---

Molecular (omic) signatures can potentially improve the precision of the clinical and pathologic sub-staging criteria currently used in routine clinical practice to quantify risk of future recurrence and death from cancer thereby better supporting patient and clinician decisions. In melanoma cancer, patients who have been diagnosed with American Joint Committee on Cancer (AJCC)^1^ stage III melanoma have diverse prognostic outcomes after resection of the lymph node metastases, with ten-year survival rates ranging from 24% to 88% in the era prior to effective systemic therapies. However, no validated molecular signatures of prognosis are currently endorsed by clinical guidelines for melanoma care, and the few that are commercially available for early stage melanoma require testing at a single location by the supplier^2^. These limitations to implementation would be greatly reduced if such signatures could be measured reproducibly on different platforms, at any suitably equipped site, and be subject to widespread external validation. We address these issues by constructing deployable low-cost assays and using a novel statistical approach to show robust application of a gene expression profile that accurately assigns prognosis in stage III melanoma across different platforms.

Previous work has shown strong associations of mRNA expression phenotypes with prognosis in stage III melanoma^3^. We identified an extended 46-gene signature that was associated with good prognosis independent of clinico-pathologic criteria, and validated it in two external datasets using microarray data^3^. The differentially expressed genes were enriched for immune activation and were subsequently shown to perform better in classifying prognosis than clinico-pathologic criteria alone, or in combination with other multi-omic (miRNA, proteome) data^4–6^. A related immune activated phenotype, detected by RNA-Seq, conferred a good prognosis in metastatic melanoma in the landmark TCGA analysis of melanoma, and was also shown to be driven by T-cell infiltrates^7^. We sought to adapt these findings (Fig. 1a) to the NanoString nCounter™ platform (NS; NanoString Technologies Inc.), based on its wide deployment and relatively low per-assay cost, and developed a robust Cross-Platform Omics Prediction (CPOP) procedure (Fig. 1b) to accurately assign risk of recurrence. The CPOP procedure enables the signature to perform consistently (by the framework shown in Supplementary Fig. 1) in transcriptome data from different gene expression platforms (microarray, RNA-Seq and NS), and in independent cohorts widely separated in time and space.

**Figure 1.**
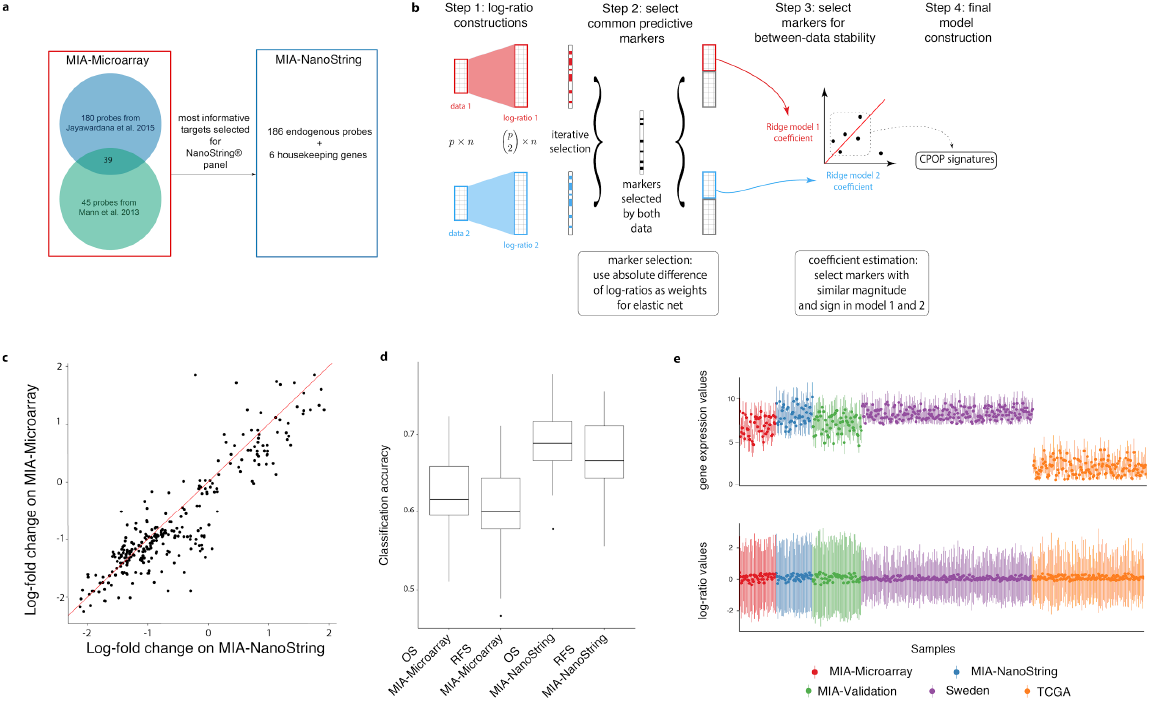
**a**, NanoString probe selection (186 probes) based on results from our previously microarray studies^3,4^. **b**, Schematic illustration of the four-steps CPOP procedure with emphasis on the stable selection of features in Steps 2 and 3. **c**, Scatter plot of log fold-change for genes common between MIA-Microarray and MIA-NanoString. **d**, Boxplot comparisons of overall accuracy for overall survival (OS) and recurrence-free survival (RFS) between the MIA-Microarray and MIA-NanoString data. The y-axis shows the classification accuracy calculated from 100 repeated 5-fold cross validation. The good/poor prognosis for overall survival (OS) and recurrence-free survival (RFS) are defined in Online Methods. **e**, Boxplot of the expression values of all genes (top panel) and all pair-wise log-ratio (bottom panel) for each sample (n = 488) in the Melanoma data collection.

We construct a signature (Fig. 1a.) consisting of 186 differentially-expressed genes that were most strongly associated with prognosis in the microarray study cohort^3,4^, together with 6 housekeeping genes (see Online Methods). This gene expression signature assay (see Supplementary Table 1 for a list of probes) was applied to melanoma biospecimens from the Melanoma Institute Australia (MIA) discovery cohort (n = 45) on both the Illumina cDNA microarray platform (MIA-Microarray) and the NanoString platform (MIA-NanoString). We further validate this signature using an independent cohort (MIA-Validation, n = 46) accrued since 2010 at the same centre, as described in Supplementary Table 2. The baseline characteristics of MIA-NanoString and the MIA-Validation cohort are described in Table 1, and Supplementary Fig. 2 shows the survival distribution of the data. We confirmed that both the gene expression (Supplementary Fig. 3) and the log-fold-differences of gene expression values between the good and poor prognosis groups measured by NanoString assay were very highly correlated (correlation = 0.9) with those originally measured in MIA-Microarray, the original discovery (Fig. 1c).

**Table 1.** Basic data summaries for five melanoma datasets, MIA-Microarray, MIA-NanoString, TCGA, Sweden and MIA–Validation. Included are the number of samples, the median age of the cohort, gender (sex), the median survival time in month and survival status. OS, RFS and DSS refers to overall survival, recurrence-free survival and disease-specific survival, respectively.

|  | MIA-Microarray<br>MIA-NanoString | TCGA | Sweden | MIA-Validation |
| --- | --- | --- | --- | --- |
| Number of samples | 45 | 139 | 210 | 46 |
| Median age (years) | 62 | 56 | 64 | 61 |
| Sex F | 17 (38%) | 59 (42%) | 86 (41%) | 14 (30%) |
| M | 28 (62%) | 80 (58%) | 124 (59%) | 32 (70%) |
| Stage/metastasis type | Stage III: 45 (100%) | Stage III: 139 (100%) | General: 23 (11%)<br>In-transit: 15 (7%)<br>Local: 11 (5%)<br>Primary: 15 (7%)<br>Regional:139 (66%)<br>NA: 7 (3%) | Stage III: 46 (100%) |
| Median survival (months) | OS: 22<br>RFS: 8 | OS: 26.9 | DSS: 17.6 | OS: 65.5<br>RFS: 9 |
| Survival status (Alive) | Yes: 19 (42%)<br>No: 26 (58%) | Yes: 78 (56%)<br>No: 61 (44%) | Yes: 108 (51%)<br>No: 102 (49%) | Yes: 26 (57%)<br>No: 20 (43%) |

A successful risk model must overcome the challenge of transferability^8^. Namely, it must be able to make predictions on independent or future data where data re-normalisation or model re-training is not practically feasible. In contrast to clinical data, gene expression data show large scale differences between datasets (see Fig. 1e) due to batch effects, intra-platform differences in protocols or other site-specific factors. Such “noise” affects the parameters estimated from the training and the validation data and thus creates undesirable inconsistencies in model performance, as illustrated in Supplementary Fig. 4.

To address these challenges, we developed a robust Cross-Platform Omics Prediction (CPOP) procedure to select transferable biomarkers (Fig. 1b, see Online Methods for a detailed description). The CPOP procedure is applicable to any type of outcome, whether diagnostic, prognostic or predictive of treatment response, and comprises four steps. CPOP first constructs features as ratios of expression of each gene compared with others, then selects those that are associated with clinical outcome. We then further select for features with high stability across datasets before fitting a final prediction model. The CPOP procedure has three strengths. First, the use of ratio-based gene expression features reduces between-data variation, as shown in Fig. 1e. Second, each feature is assigned a weight proportional to its between-data stability, in contrast to traditional modelling procedures. Finally, CPOP selects features that yield consistent estimated effects in the presence of between-data noise, thus strengthening reproducibility across datasets containing similar biological signals.

We applied the novel CPOP procedure to the MIA-Microarray and MIA-NanoString data to identify a molecular signature and corresponding prognostic model. The panels in Fig. 2b compare the hazard ratio prediction under the between-data setting (x-axis) and the within-data setting (idealistic, y-axis) and a transferable model will show high correlation of hazard ratio estimated in these two settings (see Online Methods). Here, we demonstrate CPOP’s ability to produce predictions that are on equal scale as well as demonstrating a major improvement (substantially higher correlation; 0.79 vs 0.23 in TCGA and 0.86 vs 0.06 in Sweden) compared to the commonly used Lasso regression^9^ utilised in both the external TCGA^7^ and Sweden^10^ datasets (Supplementary Fig. 5). The results of both of these analyses indicate the stability of the model’s coefficients in a between-data setting.

**Figure 2.**
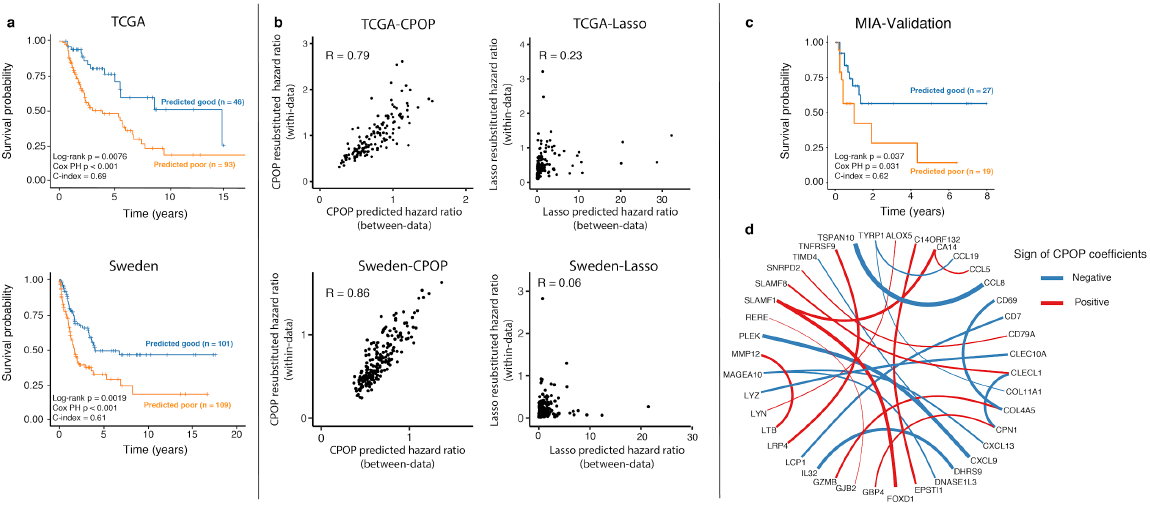
**a**, Kaplan-Meier plots show a significant difference in survival probability between the predicted good (blue line) and poor (orange line) prognostic classes on the TCGA (top panel, n = 139) and Sweden (bottom panel, n = 210). The CPOP model here is trained on MIA-Microarray and MIA-NanoString based on the RFS-defined prognosis classes. **b**, Scatter plot illustrating the concordance between between-data prediction hazard ratios and within-data (re-substituted) hazard ratios. Here we consider within-data re-substituted hazard ratios (y-axis) as the “desired or true” outcome and the cross-data prediction hazard ratios (x-axis) representing prediction across platforms without re-normalisation. We applied CPOP with a penalised Cox model with RFS as survival response for MIA-Microarray and MIA-NanoString data. R denotes the Pearson’s correlation value. **c**, Kaplan-Meier plot showing a significant difference in survival probability between the predicted good (blue line) and poor (orange line) prognostic classes on the MIA-validation data (n = 46) (including four imputed genes). The CPOP model here is trained on MIA-Microarray and MIA-NanoString. **d**, Network visualis ation of the final CPOP model highlights the ratio-based signatures developed from applying the CPOP model on the MIA-Microarray and MIA-NanoString data. Each node of the network represents a gene and an edge connecting two genes (nodes) represents the log-ratio feature that is present in the signature. The colour and thickness of each edge represents the sign and the magnitude of the size of estimated coefficients. The ratio of two genes are in alpha-numeric order.

Without further data normalisation or model modification, we assessed “model transferability” of the CPOP model on external TCGA and Swedish data and an independent MIA cohort (n = 46, see Supplementary Table 2 for detailed data description). The Kaplan-Meier^11^ plots show significant differences in survival probability between the predicted good and poor predicted classes from the CPOP model in TCGA (Fig. 2a top panel, p = 0.0015); Swedish data (Fig. 2a bottom panel, p < 0.001) and MIA-validation data (Fig. 2c, p = 0.03). As the training data (MIA-Microarray and MIA-NanoString) and the validation data (TCGA and Sweden) are based on data generated from distinct gene expression platforms, we may conclude the CPOP procedure is able to select features that are stable between platforms. Supplementary Fig. 6 shows 19 (79%) out of 24 combinations of training-testing pairs in Melanoma data showed statistically significance.

The CPOP procedure identifies the most stable relationships between gene expression values in diverse datasets, which may enhance biological interpretation. The CPOP melanoma signature is based on relative expression levels between genes and after applying the CPOP procedure onto the MIA-Microarray and MIA-NanoString data, the final CPOP signature consists of 24 log-ratio features derived from 40 genes as presented in Fig. 2d and Supplementary Table 4. In this figure, an edge connecting two genes (nodes) indicates the ratio is present in the CPOP signature and its thickness indicates the magnitude of coefficient values in the predictive model, and their direction of correlation. Genes most strongly implicated include CXCL9^12,13^,CCL8^14^, TSPAN10, PLEK, CLEC10A^15^, SLAMF1, CLECL1, MAGEA10^16^, CXCL13 and LYZ which all have been implicated as potential biomarkers in previous studies and show an enrichment in inflammatory response.

The need for model transferability and applications of CPOP is presented in another two case studies. In a case study for inflammatory bowel disease^17^, a total of 983 samples are assayed on a customised NanoString panel in three distinct batches across two years (Supplementary Fig. 7). Such a batch effect is typical of a prospective study utilising omic technologies over an extended period of time. Supplementary Fig. 8 shows that CPOP again outperforms the Lasso in concordance performance metrics and is able to make more stable cross-batch prediction on inflammation samples. Another case study is for ovarian cancer where we apply CPOP on 1,767 samples collected across ten different studies^18^ varied in clinical survival times (Supplementary Fig. 9). The heterogeneity of study cohorts presents an unique challenge that is only previously dealt using meta-analysis. We show that using a signature built from two selected studies^19,20^, we are able to make stable cross-study predictions on the remaining seven datasets (Supplementary Fig. 10), further highlighting the transferability of the CPOP signature and risk model.

The implementation of CPOP was also robust to missing data, as shown by simulation (Supplementary Fig 11). The final model implemented in CPOP is therefore highly suitable for assimilation of gene expression data from diverse platforms and cohorts. A web portal is available online (http://shiny.maths.usyd.edu.au/CPOP/) where data files of gene expression data can be uploaded and predictions for cases of stage III melanoma are obtained in real time.

In summary, in order to enable multi-centre implementation of gene expression signatures of prognosis in stage III melanoma specifically and omics signatures more widely in clinical practice we have developed a novel Cross-Platform Omics Prediction (CPOP) procedure that closes an important gap between molecular biomarker discovery and wider clinical use. The CPOP approach accounts for differences in feature scaling in omics data by performing weighted feature selection and estimation that preferentially selects stable features across multiple datasets. We validated the efficacy of the procedure at three levels. Firstly, as in-silico measures through cross-validation demonstrated high C-index and balance accuracy values. Secondly, we validated the stage III metastatic risk tool in three independent cohorts, two from publicly available data (TCGA and Sweden) and an independent in-house cohort without model modification. Finally, we demonstrate the generalisability of the procedure to two other diseases: an ovarian cancer data collection (9 data sets) and a large IBD dataset with approximate 1000 samples. Together, we deliver a molecular (omics) risk prediction platform with substantial improvements in reproducibility and stability that can be adopted in multi-centre and prospective settings.

## Methods

Methods, including statements of data availability and any associated accession codes and references, are available in the online version of the paper. The CPOP R package is available at https://sydneybiox.github.io/CPOP.

## Author contributions

JYHY and GJM conceived and funded the study. The NanoString panel was designed by EP, GP, SJS, GJM, and JYHY and the NanoString assays were established and performed by GP and VT. DS processed and performed quality checks on initial NanoString results. RAS and JFT provided the clinical and pathology input and guidance with patient clinical data curation performed by GP and VT. KW led the CPOP method development and data analysis with input from GT, SM and JYHY. KW processed and curated the data with input from JYHY. KW implemented the R package and Shiny application with help from DS and GT. DS compiled data submission to GEO with assistance from KW and GP. KW, JYHY, EP and GJM wrote the manuscript and all authors reviewed the manuscript and gave feedback. All authors read and approved the final version of the manuscript.

## Acknowledgements

The authors thank all their colleagues, particularly at The University of Sydney, School of Mathematics and Statistics, Melanoma Institute Australia, Westmead Institute and The Royal Prince Alfred Hospital for their support and intellectual engagement. The following sources of funding for each author, and for the manuscript preparation, are gratefully acknowledged: Australian Research Council Discovery Project grant (DP170100654) to JYHY, SM; Australia NHMRC Career Developmental Fellowship (APP1111338) to JYHY. Research Training Program Tuition Fee Offset and Stipend Scholarship to KW. NHMRC CRE (APP1135285) to JYHY, KW, GJM, RAS. NHMRC Fellowship (APP1141295) to RAS and NHMRC Program Grant (APP1093017) to JFT, GJM and RAS. The funding source had no role in the study design; in the collection, analysis, and interpretation of data, in the writing of the manuscript, and in the decision to submit the manuscript for publication.

## Methods

### Melanoma molecular signature assay with the NanoString nCounter platform

#### NanoString sample selection

Tumour samples were obtained from the Melanoma Institute Australia (MIA) Biospecimen Bank, a prospective collection of fresh-frozen tumours accrued with written informed patient consent and Institutional Review Board approval (Sydney South West Area Health Service institutional ethics review committee (Royal Prince Alfred Hospital (RPAH) Zone) Protocol No. X08-0155/HREC 08/RPAH/262, No. X11-0023/HREC 11/RPAH/32, and No. X07-0202/HREC/07/RPAH/30) since 1996 through MIA, formerly the Sydney Melanoma Unit. Data from two separate cohorts MIA-Nanostring and MIA-Validation are generated (see ‘Melanoma data collection’ section for details).

#### NanoString assay construction

Gene expression profiling was carried out using the NanoString nCounter^®^ platform (Seattle, WA). NanoString designed and manufactured customised probes corresponding to 192 probes. 186 of these were identified in our previously reported studies (Mann et al., 2013; Jayawardana et al., 2015). Of the 186 probes, 46 were found to be differentially expressed genes between poor and good prognosis patients (stratified based on overall survival time and vital status) (Mann et al., 2013), and 180 probes identified in a model built to predict prognosis (Jayawardana et al., 2015). Between the two studies, 39 of these probes overlapped. In instances where there was more than one probe per gene, the most informative probe based on an inclusion frequency of 30% for that particular gene was included in the NanoString panel. Six housekeeping genes were selected from a list of previously reported housekeeping genes (Eisenberg and Levanon, 2003), and had variances in the lowest quintile (20%) of the Mann et al. (2013) and Jayawardana et al. (2015) data, a differential expression p-value greater than 0.5. These housekeeping genes were selected to cover a range of high expression (CENPB, CTBP1, GNB2L1), medium (RERE, SNRPD2) and low expression (UQCR). A complete list of the customised probe set are given in Supplementary Table 1.

#### NanoString experimental procedure

Tumour RNA was extracted as previously described in Mann et al. (2013). RNA purity and concentration were assessed using Agilent TapeStation system. The nCounter^®^ gene expression assay (NanoString Technologies, Seattle, WA) was performed according to the manufacturers instructions using 100ng of total RNA. For each assay, a high-density scan (encompassing 600 fields of view) was performed.

#### NanoString hybridisation protocol

A thermal cycler is pre-heated to 67 °C. Tagset and RNA samples are removed from a freezer and thawed at room temperature. The tubes are inverted to mix and then briefly spun down. A hybridization master mix is created by adding 70 *µ*L of hybridization buffer and 7 *µ*L of the Probe A working pool directly to the tube containing the Tagset. The solution is inverted several times to mix and spun down. 7*µ*L of the Probe B working pool is added to the master mix. 8 *µ*L of master mix is added to each of the 12 strip tubes then 7 *µ*L of RNA sample is added to each tube. Strip tubes are inverted several times and flicked to increase mixing. The tubes are briefly spun and immediately placed in the pre-heated 67 °C thermal cycler, incubated for 16 hours and then ramped down to 4 °C.

#### NanoString data preprocessing and quality assessment

NanoString data was read into `R` using the `NanoStringQCPro` package (Nickles et al., 2019). For the purpose of illustrate cross-platform noise, we performed a simple log 2 transformation on the raw counts. This allow us to assess if the panel can facilitate prospective experiment without model modification. Of all the samples measured in the “MIA-NanoString” data, 45 samples overlap with a previous microarray study (Mann et al., 2013; Jayawardana et al., 2015), named “MIA-Microarray”. This allows us to correlate genes between the two data in Supplementary Fig. 3. Of the 192 common genes (matched through official gene symbols), the median correlation is 0.86 with the first and third quartiles being 0.79 and 0.90 respectively.

#### Statistical analysis - RFS assessment

Clinical follow-up time for samples presented in (Mann et al., 2013) and (Jayawardana et al., 2015) is updated in 2018 and we define recurrence-free survival (RFS) as the time difference between the date of first recurrence after tissue banking and the date of tumour banked. Based on RFS, we further defined:

- “Good” prognosis group being RFS greater than 4 years and alive with no sign of recurrence.
- “Poor” prognosis group being RFS less than 1 year and died due to melanoma.

This resulted in 19 samples in the good prognosis group and 26 samples in the poor prognosis group. Using this RFS-defined prognosis classes for these 45 samples, we use limma (Ritchie et al., 2015) to compute moderated t-statistics for all 192 overlapped genes.

#### Statistical analysis - Prognostic assessment

We validate the prognostic assessment for the MIA-NanoString datasets by comparing its performance accuracy for both overall survival (OS) and RFS against the same samples in the MIA-Microarray data. Performance accuracy is obtained by using the `ClassifyR` (Strbenac et al., 2015) package to compute 100 repeats of 5-fold cross-validation with the limma (Ritchie et al., 2015) feature selection and support vector machine classifier. This is specified via the parameters `ResubstituteParams(nFeatures = c(20, 50, 100), performanceType = “balanced error”, better = “lower”)` before executing the function `runTests`. All analysis are performed in `R` (R Core Team, 2019) version 3.6.2.

#### Melanoma data collection

We describe five melanoma datasets mostly consisting of late-stage samples, see Supplementary Table 2. Three of these data are samples from the MIA and the other two are publicly available data. Of the three MIA data, one is measured by Illumina microarray technology and two are measured by our customised NanoString assay as described in the ‘Melanoma molecular signature assay’ section.

1. **MIA-Microarray**: A published gene expression study (Illumina platform) with 45 stage III melanoma subjects from Melanoma Institute Australia.
2. **MIA-NanoString**: An in-house gene expression data constructed in 2018 using the customised NanoString assay. The 45 samples are the same as the MIA-Microarray cohort described above.
3. **MIA-Validation** This independent validation cohort contains 46 samples that are age-matched and have similar characteristics to the MIA - NanoString were used to validated the CPOP procedure. Due to manufacturer error, 12 genes are unavailable to be run in this cohort and treated as missing.
4. **TCGA**: We downloaded the RNA-Seq data consisted of 472 samples from Network (2015) on 28th July 2017 using the `TCGABiolinks` package (Colaprico et al., 2016) in R (R Core Team, 2019) and Bioconductor (Gentleman et al., 2004). We processed the data into log2-FPKM values and only retained 458 samples with survival times and status recorded. Out of these samples, 169 samples are labelled as Stage III, and after removing 30 samples that MIA has contributed to, we are left with a total of 139 independent samples to be used in the evaluation of survival analysis.
5. **Sweden**: A published microarray study in (Cirenajwis et al., 2015). We retain all 210 samples for survival-based evaluations as there is a lack of cancer staging information in the data. Processed data was downloaded on 12 January 2020 via the `GEOquery` package, Davis and Meltzer (2007).

#### Ovarian cancer data collection

The curation of this data collection closely follows from Waldron et al. (2014) and the analysis pipeline from Yoshihara et al. (2012). All data are downloaded through the `cureatedOvarianCancer` Bioconductor package (Ganzfried et al., 2013). Out of the ten datasets in Table 2 of Waldron et al. (2014), we make some key modifications to the selection of data:

- leaving out Konstantinopoulos et al. (2010) data for its incomplete sample annotations;
- leaving out Dressman et al. (2007) data as this article was retracted;
- swapping the TCGA microarray data (Bell et al., 2011) for the RNA-Seq data; and
- adding one extra data, abbreviated as “Japan B” from Yoshihara et al. (2012).

We focus on the 126 gene signature reported in Yoshihara et al. (2012) and select genes that are also present in all nine datasets. This results in 94 genes with corresponding 4,371 log-ratio features. This ovarian data collection from nine different datasets consists of samples described in Supplementary Table 3.

#### Inflammatory bowel disease data

This data is measured on a NanoString platform with genes originally selected to study disease-associated risk loci in inflammatory bowel disease (IBD), (Peloquin et al., 2016). We try to classify all 983 samples as either inflamed or not inflamed learning from 712 genes. The original authors provided the raw NanoString data on Gene Expression Omnibus repository under the accession number GSE73094. We perform log2-transformation on the raw counts data. This IBD data is chosen as the experiment extends over a few years with obvious batch effect as the chemical reagent was changed twice. This change in the use of reagent creates three batches, IBD2 (*n* = 303), IBD3 (n = 295) and IBD4 (*n* = 385) that mimics the implementation challenge such prognostic (or risk) models will face when it is implemented in a prospective setting. Supplementary Fig. 7 shows the batch effect in this IBD via a sample boxplot and a principal component analysis (PCA) plot. Previous effects in addressing the stability in data quality is through normalisation techniques, for example, in Molania et al. (2019). CPOP distinguishes itself from normalisation techniques through the use of log-ratios and making predictions simultaneously.

#### Cross-Platform Omics Prediction (CPOP) Overview

Cross-Platform Omics Prediction (CPOP) is a procedure that enables sample prediction across gene expression datasets with different scales (e.g. different sample means). We will use the generic phrase of “scale difference” to encompass all situations where multiple gene expression data exhibit different scales in the data due to, for example, the use of different experimental instruments/platforms or drifts in measurements in a prospective setting. We use the term ‘biomarker’ and ‘feature’ interchangeably. We will use the term *predictive* in a statistical sense and the term *predictive markers* in a generic way referring to all forms of biomarkers whether they are diagnostic, prognostic or predictive. We use the term *training set* interchangeably with *reference set* (or *sets*), and restrict usage of the term *test set* or *validation set* to situations with known patient outcome i.e. to situations where we are assessing or comparing the performance of CPOP. We use the term *test sample* or *validation sample* when the unknown subjects are to be predicted.

A major consideration in developing CPOP is to make prediction on a single-sample without normalisation or combining it with additional data. The CPOP procedure has the following three key characteristics:

1. CPOP uses (log)-ratios of genes as biomarkers (features), which are more stable than using individual gene expression values (see Step 1 of CPOP).
2. CPOP uses Elastic Net models to perform feature selection using weights proportional to the stability of features across more than one data set. This allows the selection of common predictive markers (see Step 2 of CPOP).
3. CPOP selects for features with high similarity in their between-data estimated effects (Step 3 of CPOP).

### CPOP - four-step procedure

#### Statistical background

Suppose we have a gene expression data matrix ***X*** ∈ ℝ^*n× p*^ where *n* is the number of samples and *p* is the number of genes on a gene expression platform. We define the “log-ratio matrix” as a matrix ***Z*** of dimension ℝ ^*n*×*q*^ where 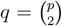 and each column of ***Z*** is the pairwise difference between two log-transformed columns in ***X***. Formally, each column of ***Z*** is given by enumerating all log-ratio features log(***x***_*l*_) log(***x***_*m*_) for 1 ≤ l < m ≤ p. Thus, each feature in the ***Z*** matrix is the log-ratio of the expression values of two genes. For the given log-ratio matrix ***Z*** ℝ ^*n*×*q*^, we denote each row of the matrix as ***z***_*i*_ for patient i = 1,. .., n. Let ***y*** ∈ ℝ^*n*^ be a vector that measures each patient’s clinical outcome (e.g. patient tumour shrinkage in millimetres or prognostic outcome).

The **weighted Elastic Net (WEN) model** is a regularised regression model that solves for a regression coefficient vector 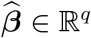 with the optimisation equation:

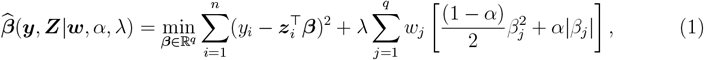

where, *λ* ∈ (0, *∞*] and *α* ∈ [0, 1] are tuning parameters and ***w*** = (*w*_1_,. .., *w*_*q*_) is a sequence of weights placed on each of the q features. This will be explained in context of our cross-platform prediction later. The *first component* of Equation (1) is a linear loss function that measures the difference between the fitted value 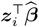 and the observed response value *y*_*i*_ for sample *i*. This component can be readily substituted with any appropriate non-linear loss function depending on the variable type of the response variables (e.g. in ‘Performance evaluation’ section, logistic and Cox models are used to deal with binary and survival responses, respectively). The *second component* of the equation is a penalty on the magnitude of the estimated regression coefficients 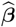. This component mixes a *L*_1_-norm penalty and a *L*_2_-norm penalty through the use of the *α* tuning parameter. The , *λ* tuning parameter controls the total strength of penalisation in the overall equation.

### CPOP procedure

This procedure is designed to handle both cross-platform data and prospective implementation data over time since the differences in data scale affect both of these data settings. We describe the **CPOP procedure** below:

Step 1. **Log-ratio matrices construction**: Suppose we have two gene expression data and the associated log-ratio matrices as 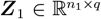 and 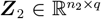, where *n*_1_ and *n*_2_ are the samples sizes for the two datasets. We do not impose the restriction of paired samples across the two data, however, we assume the two data measure the same q log-ratio features or we restrict our modelling to the common q log-ratio features between two datasets. For both data, we also have a clinical outcome measurement, denoted as 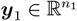 and 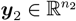 associated with data 1 and 2 respectively.

Step 2. **Selecting common predictive features**: compute a sequence of non-negative weights w_*j*_,j = 1,. .., q that measure the column-wise statistical concordance between ***Z***_1_ and ***Z***_2_. Fit a WEN model for both (***Z***_1_, ***y***_1_) and (***Z***_2_, ***y***_2_) using w_*j*_,*j* = 1,. .., *q* to obtain estimated regression coefficients 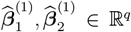 with a penalty parameter *α* ∈ (0, 1]. Note the superscript denotes these regression coefficients are in the first step of CPOP.

Since WEN generates sparse estimates for *α* ≠0, thus it also naturally selects features from our data as those features with non-zero estimates in 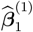 and 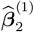. Define a feature set 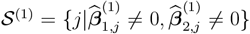 that collects all non-zero features selected into both models in both data.

In this paper, we primarily focus on the use of mean-difference weights: w_*j*_ = |mean(***Z***_1*j*_) *-* mean(***Z***_2*j*_) | for each *j* = 1,. .., q, whereas other choices are also available in our `CPOP` package.

Step 3. **Selecting features with between-data stability**: define 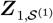 and 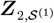 as the matrices that we obtain when subsetting ***Z***_1_ and ***Z***_2_ to only the features present in 𝒮 ^(1)^. Then, fit an unweighted ridge regression model (i.e. a WEN model with no weights and *α* = 0) onto 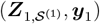 and 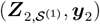 to obtain 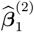 and 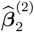. Define another feature set 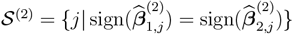.

We also include an additional step by iteratively fitting ridge regression to both 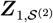 and 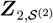 and in each iteration, we update the feature set 𝒮 ^(2)^ by removing all features that do not satisfy 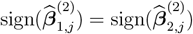. This removal of features using the ridge models means that the size of 𝒮 ^(2)^ is non-increasing with each iteration. This iterative step terminates when there is no further reduction in the size of 𝒮 ^(2)^.

Step 4. **Final model estimation**: the final CPOP models are the unweighted ridge regression models fitted onto 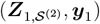 and 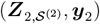. We will refer to these models as 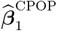 and 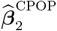, respectively. Predictions on new samples could be made by using the coefficients 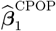 or 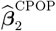 or taking the average of the two to produce a singular 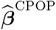.

### Practical implementation

#### Imputation

One particular important issue is the handling of missing values. CPOP model training assumes two complete data. However, if certain genes cannot be measured in a test/validation data, then imputation on the gene-level data (***X***_test_) will be necessary prior to the calculation of the log-ratio matrix ***Z***_test_. We make no particular recommendation on what imputation method to use as there are many good methods and preferences as well as their vary among practitioners. However, we highlight a special case of missingness in gene expression data, which is when a gene is not measured at all. This situation arises typically when a gene fails quality control and any numerical values are deemed as invalid. In this case, we propose to use the non-missing gene values in the two gene-level training data of CPOP (***X***_1_ and ***X***_2_) to impute on ***X***_test_. While a variety of methods can be used, in the `CPOP` package we provide a function that utilises the Lasso estimator from the `glmnet` package (Tibshirani, 1996; Friedman et al., 2010) to make this imputation.

#### Setting a CPOP prediction cut-off

Applying the CPOP model to any new sample of interest will produce a predicted value similar to that of a linear regression model. In order to produce a binary class prediction (e.g. good vs poor prognosis), a single cut-off independent of between-data scale differences is needed. Here, we will set a cut-off at 0 for the linear predicted response variable 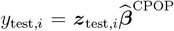 to produce binary class predictions. This cut-off is chosen for its ease of interpretation in the context of two common types of clinical variable modelling. In binary classification, a cut-off for this linear predicted response implies a cut-off at 0.5 for the predicted probability, see section on ‘Performance evaluation’. This is a threshold which a sample is equally likely to be assigned to one of two binary classes. Similarly, in Cox regression model in survival analysis, this cut-off at 0 corresponds to a cut-off at 1 for the predicted hazard ratio, see section on ‘Performance evaluation’. Here, a value great than 1 implies an at-risk sample. Though it should be noted, as with all statistical models, this cut-off is only sensible if the validation data is biologically and clinically similar to training data.

This notion of a data-independent cut-off is closely related to the challenge of between-data scale differences and forms a critical part in our evaluation. A data-adaptive choice of this cut-off on the linear prediction values, for example, using the median, could easily bias the result in assuming there is about half of high-risk incoming samples, a poor assumption in prospective testing. On the other hand, the popular area under the receiver operating characteristic curve (AUC-ROC, Fawcett (2006)) metric for binary classification and the survival concordance index (C-index, Harrell et al. (1996)) are not appropriate measures as they avoid making a cut-off and thus mask the scale differences in the data. Nonetheless, we choose to report on both of these metrics in this manuscript so comparisons may be made with other publications.

#### Performance evaluation

We propose an evaluation framework with an emphasis on producing between-data predictions that are robust to scale differences. Supplementary Fig. 1 summarises the evaluation framework we have. For the evaluations below, we choose to use the prediction values averaged between the two ridge models at Step 4 of the CPOP procedure. Here, we choose to focus on two most common response data seen in clinical studies:

- Binary classification response variable (e.g. good vs poor prognosis) can be modelled using the (penalised) logistic regression loss function. In logistic regression, the probability for assigning a sample i can be written as p_test,*i*_ = 1/[1 + exp(*-*y_test,*i*_)].
- Survival time response variable (e.g. recurrence-free survival) can be modelled using (penalised) Cox proportional hazard loss function. In Cox proportional hazard model, the hazard ratio can be written as 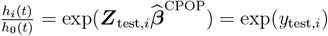.

Both can be modeled using the `glmnet` package (version 3.0-2) (Friedman et al., 2010).

### Evaluation Metrics and settings

We consider three broad classes of performance metrics to capture survival performance, classification performance and concordance performance. Supplementary Fig. 1 summarises the various metrics we use in connection with the CPOP training and testing data. In our evaluation setting, majority of the metrics can be calculated under 100 repeated 5-fold cross validation.

### Survival performance metrics

Where survival time is available in the test data:

1. **C-index**: we take the predicted values from either a CPOP or a Lasso model and fit a classical Cox regression model (i.e. without penalisation) together with age and gender. The C-index Harrell et al. (1996) of this Cox model is reported. The C-index is defined as:

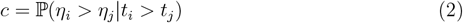

where t_*i*_ and t_*j*_ are survival times of sample *i* and *j* respectively and *η*_*i*_ and *η*_*j*_ are linear predicted values in the classical Cox regression model. Survival analysis is performed using the `survival` package (Therneau, 2020).
2. **KM-plot**: we create a binary split of predicted samples at the hazard ratio of 1. This binary split means we can construct a Kaplan-Meier (Kaplan and Meier, 1958) survival plot (KM-plot) using the `survminer` package (Kassambara et al., 2020). The log-rank statistic associated with this KM-plot is also reported.
3. **Log-rank test p-value**: similar to the KM-plot evaluation above, we also compute the log-rank test p-value of the binary split of predicted sample classes.

**Concordance performance metrics**. We propose two additional statistics to measure between-data concordance. For a validation data, we may calculate

- a re-substituted value, 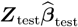 and use this value as the “gold-standard” against
- a prediction value 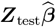.

In the application to Melanoma data collection, we evaluate CPOP against the competing Lasso model, where we use a Lasso-Cox model (results shown in Fig. 2b) for feature selection and then these features are then fitted using a ridge-Cox regression model. That is, we choose 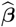 to be the ridge regression coefficients, 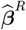, fitted using only features selected CPOP or the Lasso. Denoting 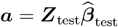 and 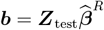 . This procedure ensures that we can make fair comparisons across two distinct feature selection methodologies.Thus, two statistics we propose are:

1. **Pearson’s correlation** between ***a*** and ***b***, which measures the concordance between the between-data prediction and the within-data re-substitution value. A higher positive value implies a higher quality of between-data prediction. To visualise this evaluation, the CPOP models and the Lasso-Cox models are placed on the *x*-axes of Fig. 2b and the re-substituted prediction values are placed on the *y*-axes and the Pearson’s correlation is calculated.
2. **Identity distance** between ***a*** and ***b***, which is defined as:

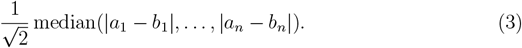 This metric is based on the simple geometric fact that for a given point (a, b) *∈* ℝ^2^, the quantity 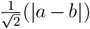 measures the perpendicular distance between the point to the identity line *y* = *x*. Hence, this identity distance can measure the average deviation between the within-data resubstitution values (***a***) and the between-data prediction values (***b***). A lower value implies a better agreement between the two quantities.

**Classification performance metrics**. For cases where we are interested in binary classification (e.g. good vs poor prognosis) and all metrics below can be repeated applied under the 100 repeated 5-fold cross validation setting.

1. **AUC-ROC**: we use the `yardstick` package (Kuhn and Vaughan, 2020) to compute the AUC-ROC. Though it should be noted that this is not the most appropriate metric, its construction masks the effect of data scale differences.
2. We create a binary split of predicted samples at the predicted probability of 0.5 (e.g. predicted good prognosis class vs predicted poor prognosis class). The predicted class and true class labels defined for each data makes up a confusion matrix with four categories: true positives (TP), true negatives (TN), false positives (FP) and false negatives (FN). The predicted binary class are evaluated using classical classification statistics and computed using the `yardstick` package (Kuhn and Vaughan, 2020):
  a. **Precision** of prediction, defined as:

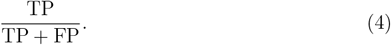
  b. **Recall** of prediction, defined as:

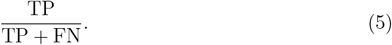
  c. **Balanced accuracy**, defined as:

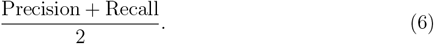
  d. **The Matthews correlation coefficient** (MCC), defined as:

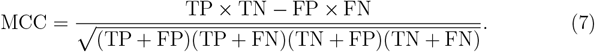
  e. *F*_1_ **statistic**, defined as:

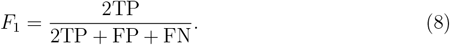

All of these metrics range from 0 to 1. In the application to Melanoma data collection (results shown in Fig. 2c), we use the CPOP model to make prediction on the MIA-validation 46 samples (reduced from five batches each consist of 12 samples by removing technical replicates).

### Evaluation on ovarian data collection

We apply the CPOP procedure with a penalised Cox loss function on the Japan A (Yoshihara et al., 2012) and Tothill (Tothill et al., 2008) data as the dual training set ^1^. This collection of ovarian data is quite heterogeneous (Supplementary Fig. 9). Through careful data curation, we selected a subset of data where the range of survival times overlap between multiple data, however, the degree of separation by survival status still varies from data to data. Thus, this heterogeneity still impacts our analytics.

### Evaluation on inflammatory bowel disease data

Treating IBD2 and IBD3 as the training set, we apply the CPOP procedure with a penalised logistic loss function on the inflammation status of the samples. The selected features are then refitted back on IBD2 and IDB3 separately using ridge regression. These ridge regression models are then used to predict on the inflammation status of samples in IBD4 (Supplementary Fig. 12).

## Footnotes

1 Due to instability in the coefficient estimates in the second step of CPOP, we use an alternative approach of retaining features with estimated coefficients within 0.5 units with each other.

