## Supplementary Materials for "Cross-Platform Omics Prediction procedure: a game changer for implementing precision medicine in patients with stage-III melanoma"

### Supplementary Figures

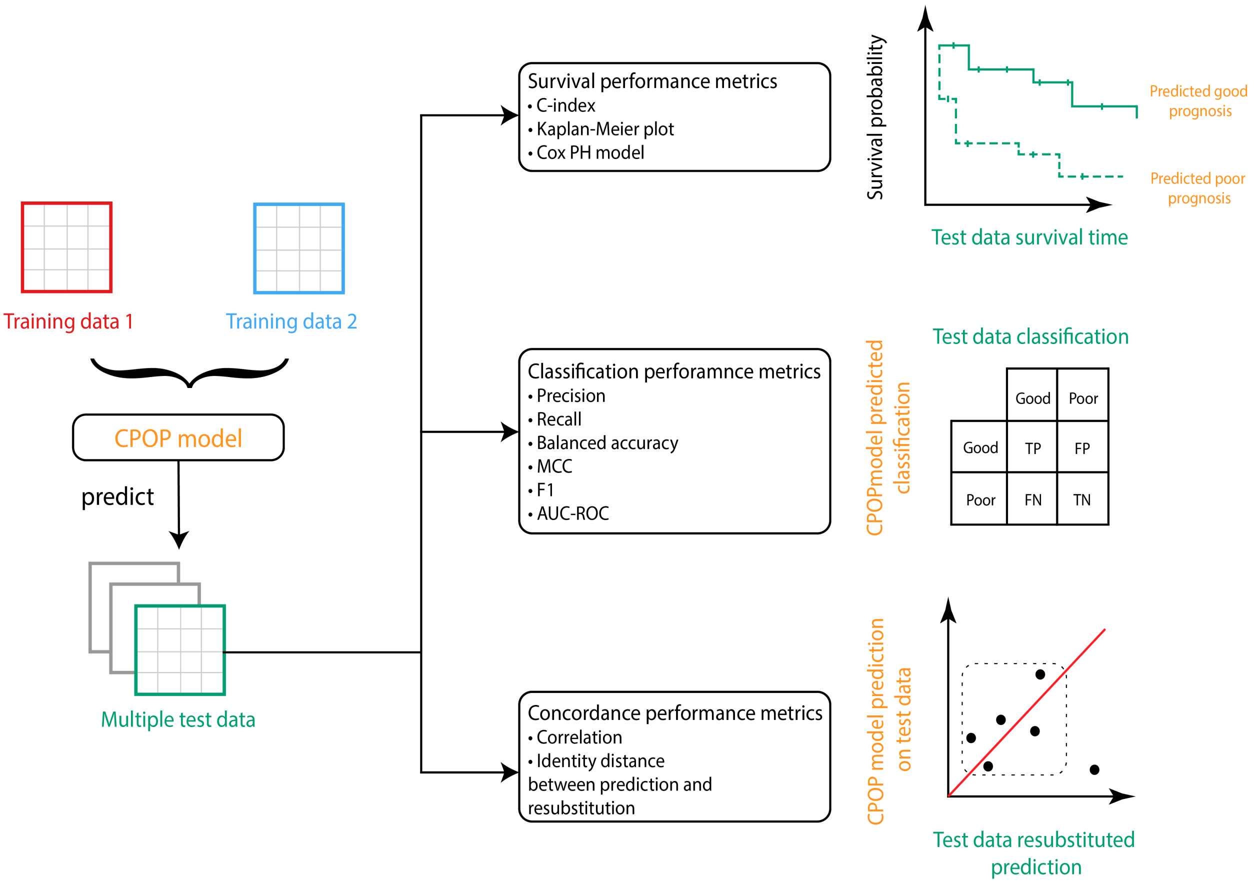


Supplementary Figure 1: Schematic illustration of the evaluation framework for CPOP using three different classes of metrics.

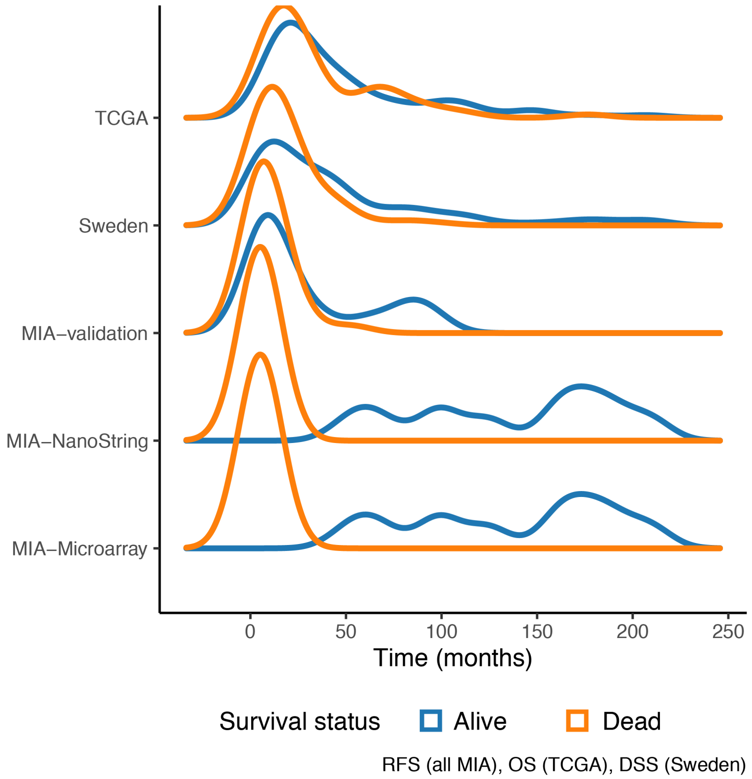


Supplementary Figure 2: Distribution of the survival time for the melanoma data collection, stratified by survival status. All data illustrate here use disease-specific survival times or recurrence-free survival except for TCGA data where overall survival is used.

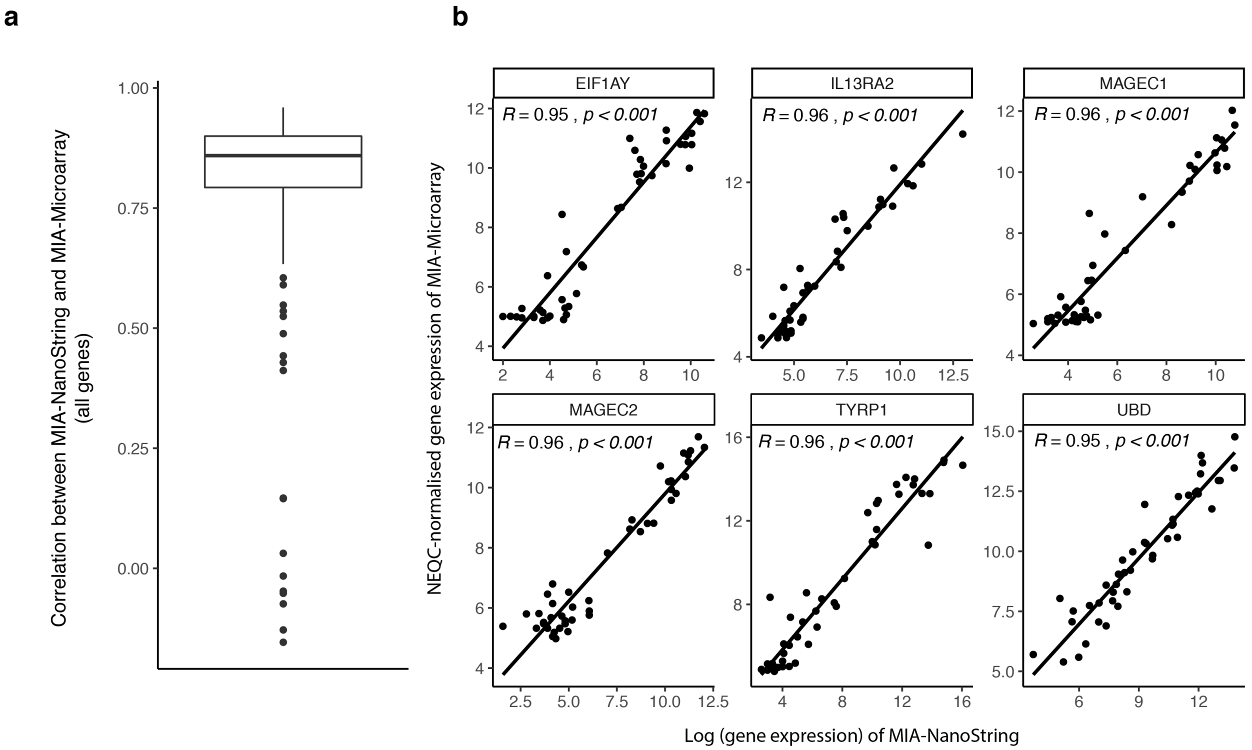


Supplementary Figure 3: **a**, Boxplot of Pearson's correlation of gene expression values between MIA-Microarray and MIA-NanoString for 192 common genes. **b**, Scatter-plot of six genes with the highest correlation across the two data.

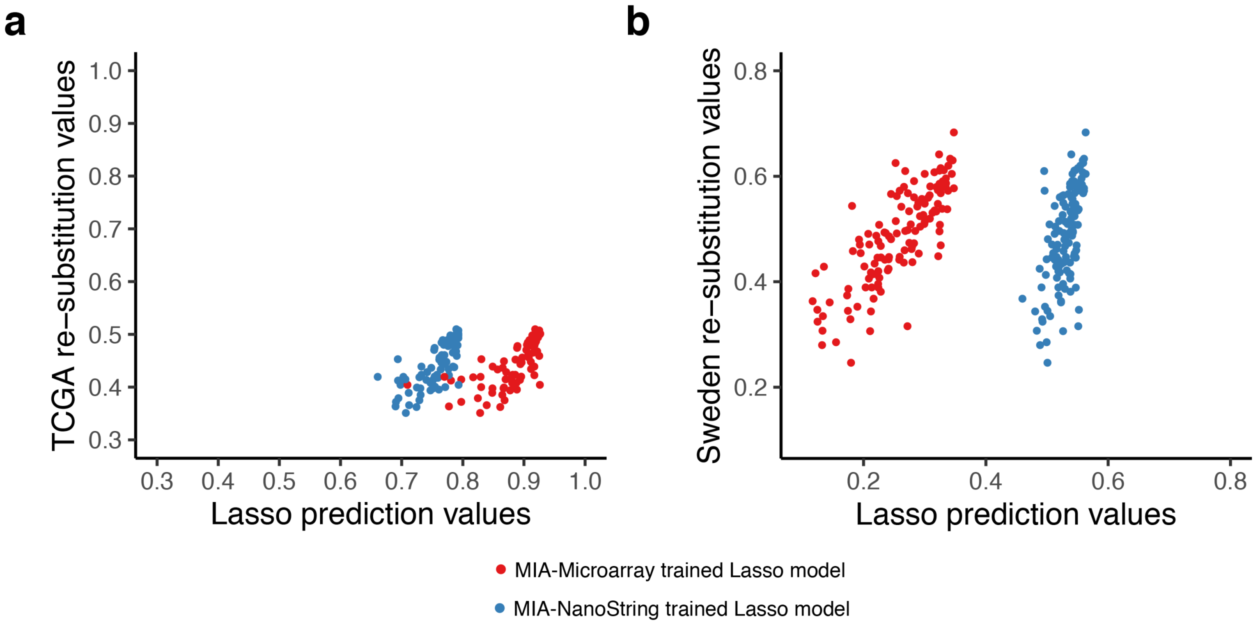


Supplementary Figure 4: This plot highlights the statistical challenges of Lasso regression, where scale difference in the data can lead to scale difference in the prediction which will affect the translational interpretation. On the x-axis, we plot the prediction values from a Lasso model trained on MIA-Microarray (red points) and another trained on MIA-Nanostring (blue points), with both models making predictions on the TCGA (panel **a**) and Sweden (panel **b**) samples. One the y-axis, we plot the re-substituted Lasso models where we trained and tested on the TCGA data (panel **a**) and Sweden (panel **b**). As the re-substituted values (y-axis) are identical between-data platform, we can then compare the scale of prediction values between the MIA-Microarray-trained model and the MIA-NanoString-trained model (x-axis) and see an obvious scale difference between the two. Furthermore, both of these between-data prediction values are on a very different scales to the re-substituted ("gold-standard") values from TCGA and Sweden.

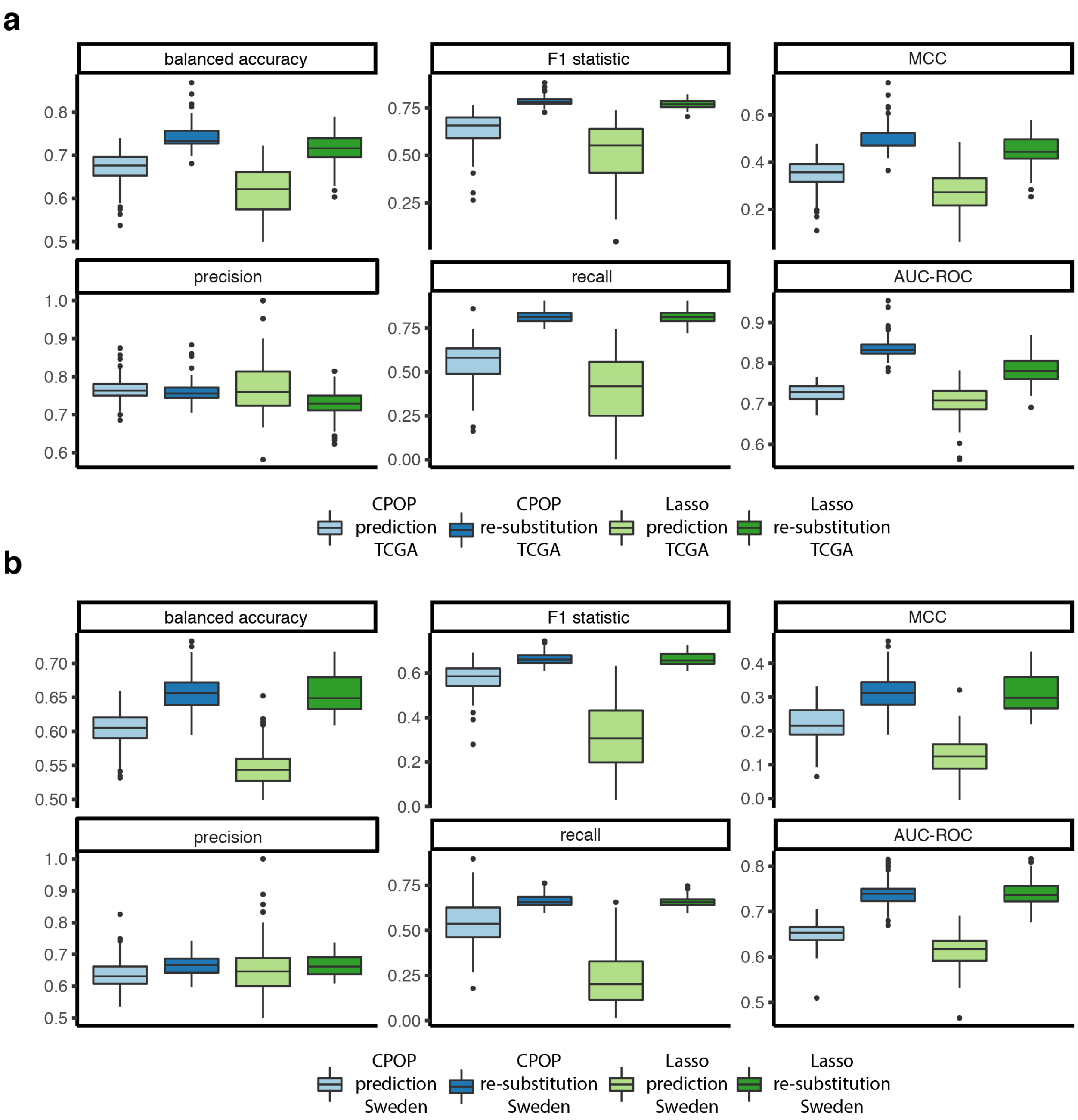


Supplementary Figure 5: We perform 20 repeated 5-fold cross-validation to select 100 different sets of features where each CPOP and Lasso feature set is constructed from 80% of MIA-Microarray and MIA-NanoString samples. We then plot the classification performance metrics from Supplementary Fig. 1 here. We compare the CPOP and Lasso predicted probabilities in panel a and b respectively. We also add the within-data re-substituted values for CPOP and Lasso for reference. While within-data performance of CPOP and Lasso are similar, CPOP consistently perform better than Lasso in terms of between-data performance.

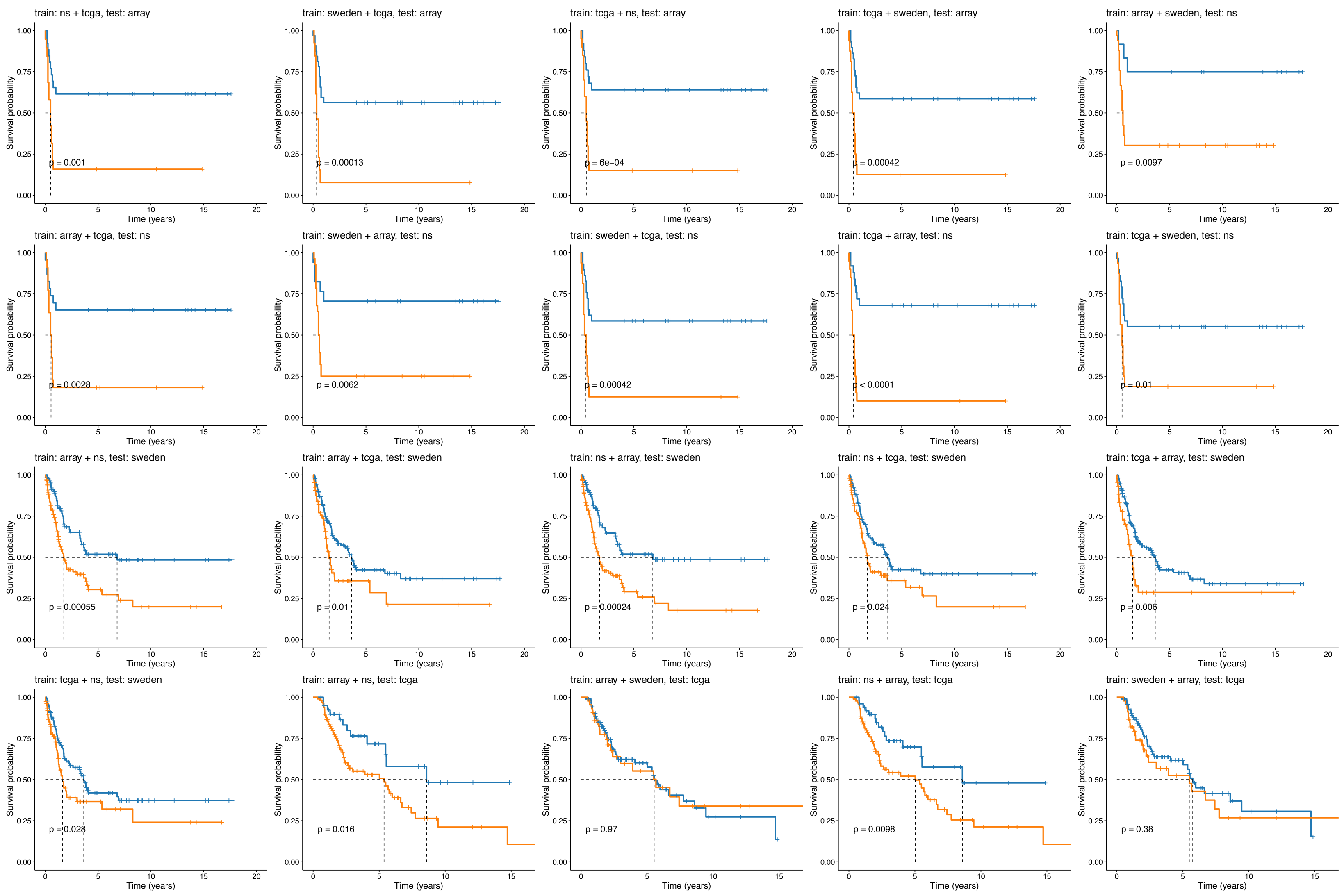


Supplementary Figure 6: Assessment of melanoma data with survival performance metrics. Kaplan-Meier plots show a significant difference between the predicted good (blue line) and poor (orange line) prognostic classes on different training-testing pairs from four of the melanoma data collection (MIA-Microarray, MIA-NanoString, TCGA and Sweden). Here, we show that the general applicability of the CPOP procedure through KM-plots of all 24 combinations of training-testing set of the MIA-Microarray, MIA-NanoString, TCGA and Sweden data. While not every training and testing combination shows a statistical difference, a majority (19 out of 24, or 79%) of pairs did.

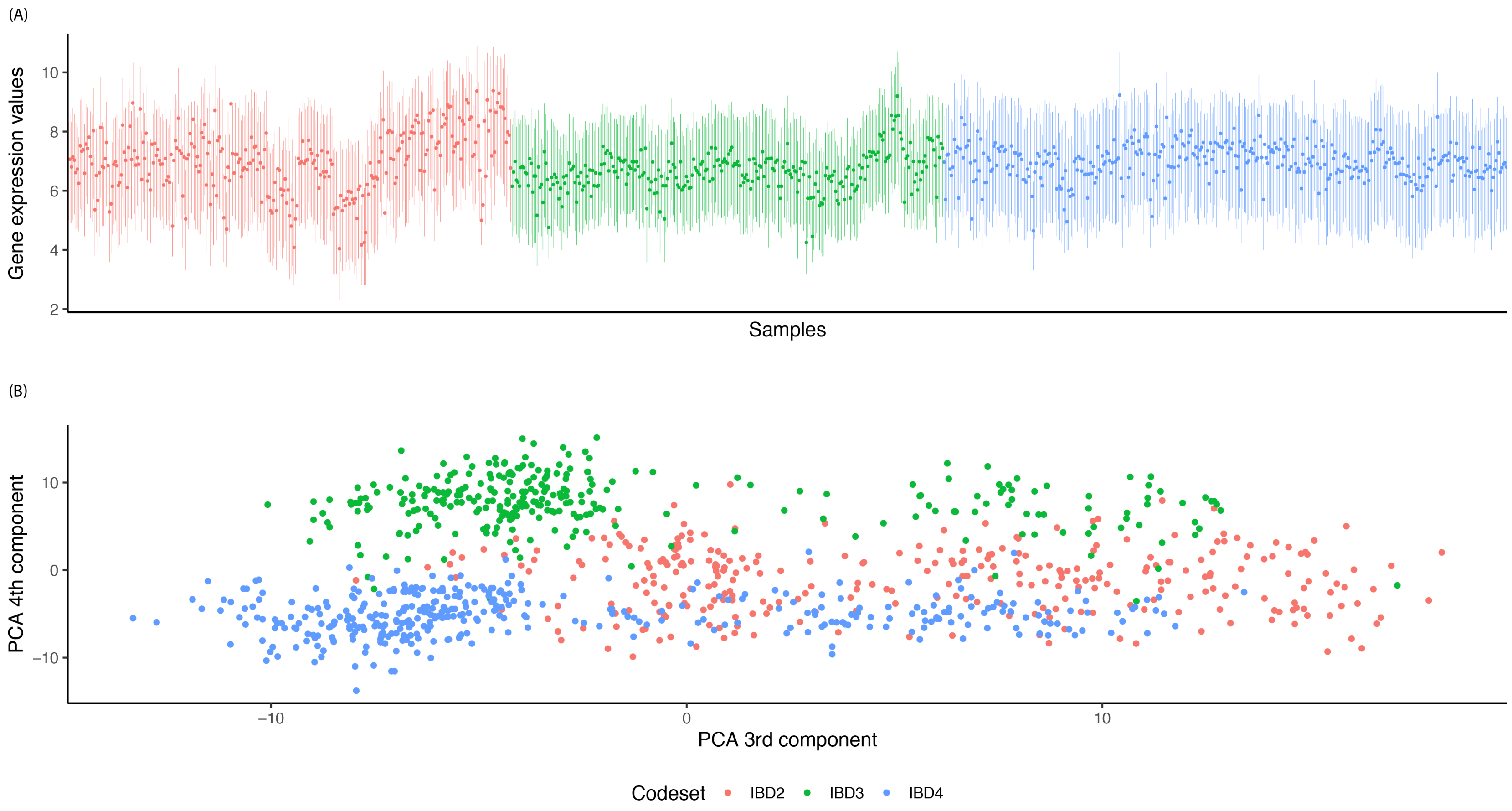


Supplementary Figure 7: (A) Boxplot of gene expression values for the IBD data, coloured by the reagent codeset. (B) PCA plot using the 3rd and 4th components, coloured by the reagent codeset.

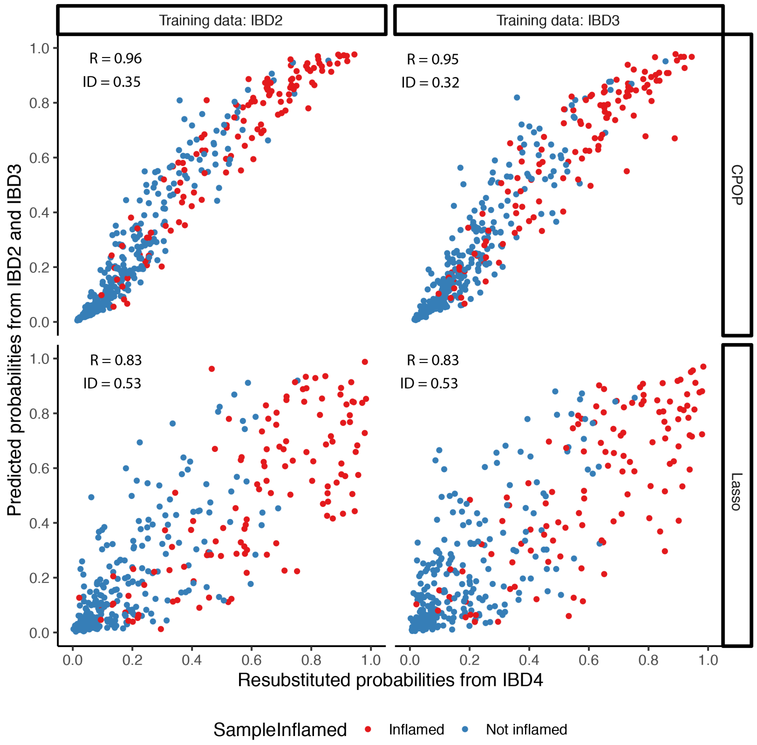


Supplementary Figure 8: Comparing the within-batch re-substituted probability and the between-batch estimated probability by CPOP and Lasso on the inflammatory bowel disease data. Here, the x-axis represents re-substituted model risk from the IBD4 batch, the y-axis represents between-batch predicted risk from the IBD2 and IBD3 on the IBD4 batch. R denotes the Pearson's correlation value and ID denotes the identity distance value. Both the CPOP and Lasso method achieve similar binary classification performance (e.g. balanced accuracy are at 0.75 and 0.78, respectively). However, CPOP outperforms the Lasso in concordance performance metrics, with CPOP showing a tighter clustering around the identity line than the Lasso prediction, as shown via the Pearson's correlation values and identity distances, illustrating the potential of the CPOP model to avoid re-normalisation over time.

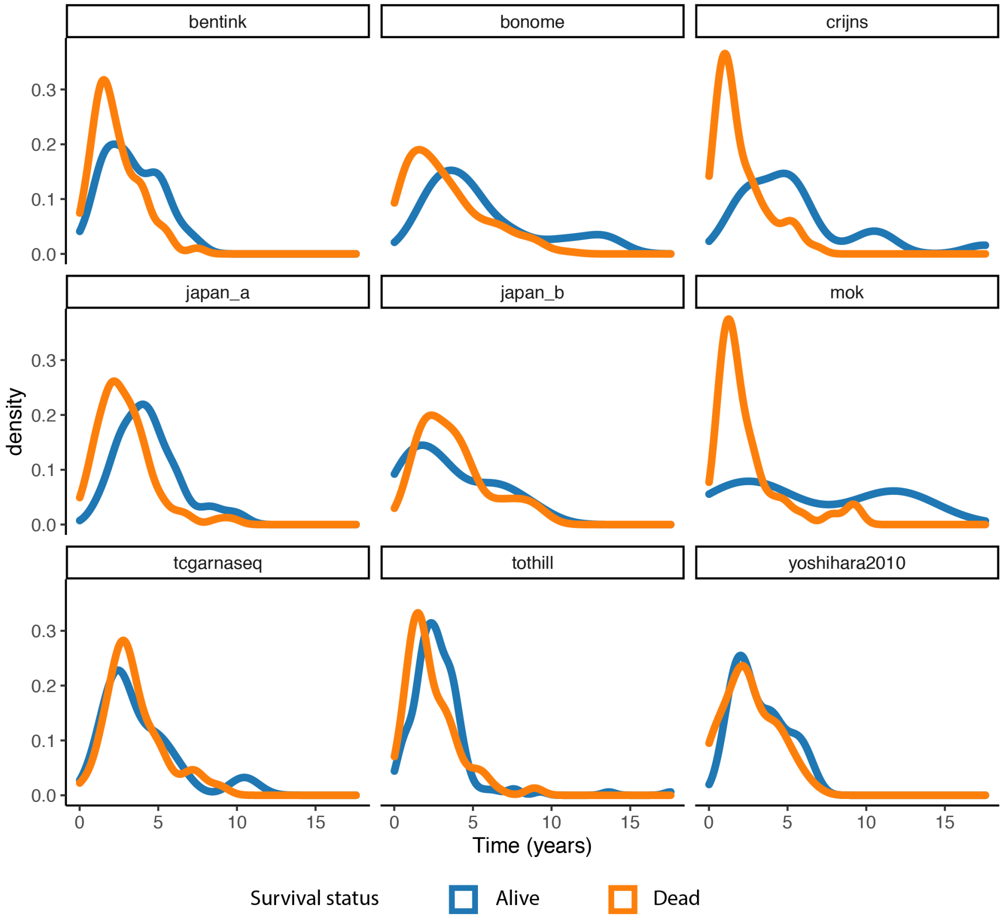


Supplementary Figure 9: Distribution of the survival time for the Ovarian cancer data collection, stratified by survival status. All data illustrate here uses overall survival time.

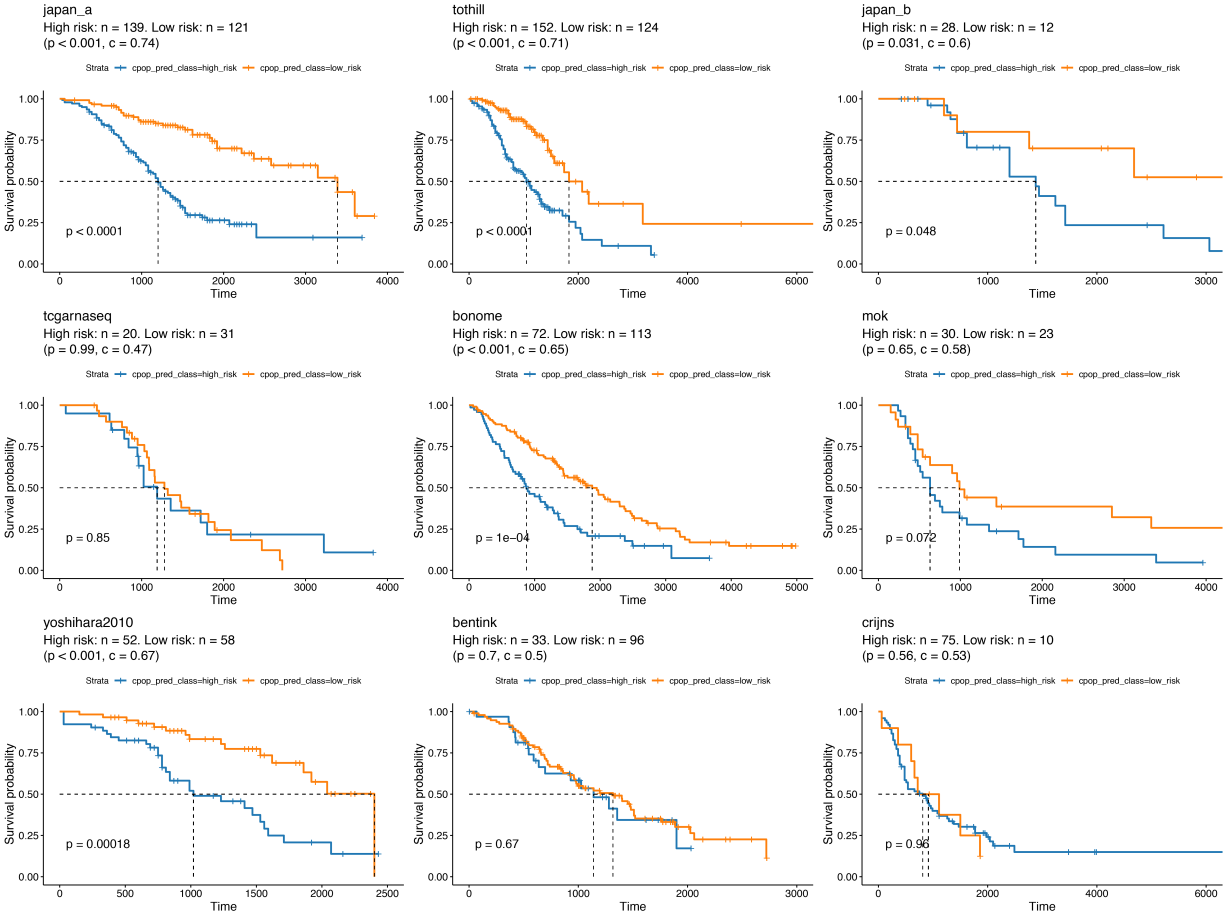


Supplementary Figure 10: Assessment of the ovarian data with survival performance metrics. Kaplan-Meier plot of CPOP prediction on all the nine ovarian data showing the survival probability between the predicted good (blue line) and poor (orange line) prognostic classes. We build the model using the CPOP procedure with a penalised Cox loss function on the Japan A (Yoshihara et al., 2012) and Tothill (Tothill et al., 2008) data. Five out of the nine data show a statistical significance between good and poor predicted outcomes. This illustrates the high predictive strength of the CPOP method. While not all data achieved statistical significance as the original publication (Waldron et al., 2014), it should be noted that $z$-score standardisation (standardisation with mean equal to 0 and variance equial to 1) is applied on all data prior to modelling in (Waldron et al., 2014). On the other hand, our CPOP evaluation avoids this transformation for the reason that we wish to best evaluate our method in a single-sample prediction situation where $z$-transformation is not impossible.

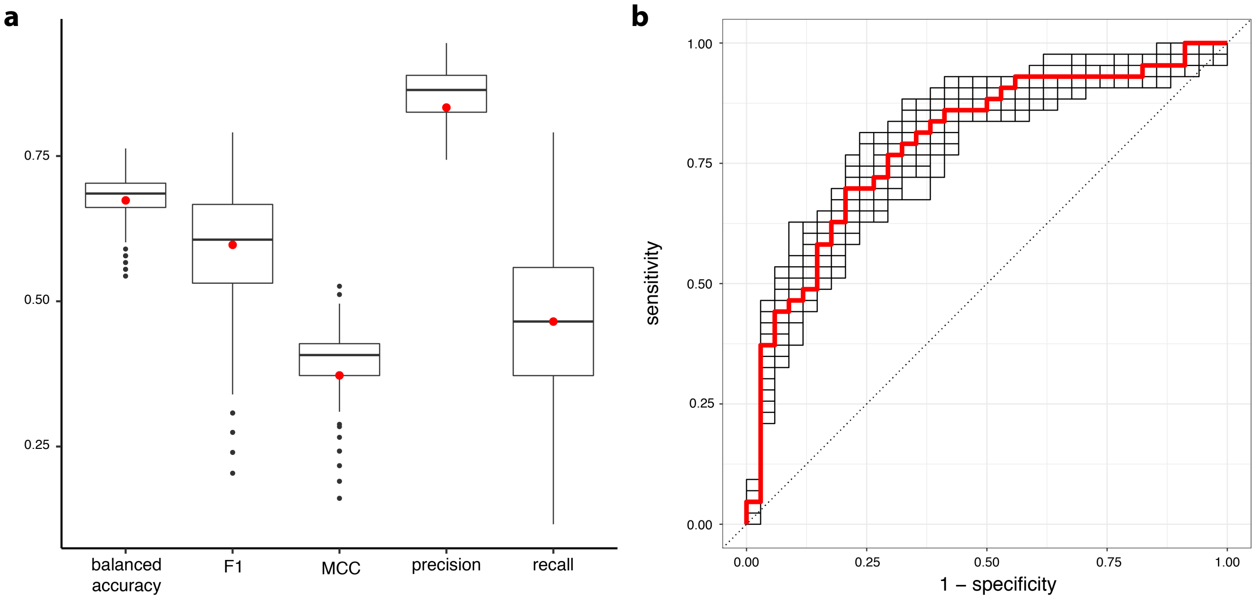


Supplementary Figure 11: Assessment of prediction model with missing values. We introduce missing values into the TCGA data by randomly removing 5 genes and use the training model to impute the missing genes before calculating the log-ratio matrices and corresponding CPOP prediction. The process randomly removing of 5 genes is repeated 100 times. Panel **a** illustrates the prediction performance using five difference performance metrics (balance accuracy, F1, MCC, precision and recall). The red points represent the prediction performance on the complete TCGA without any missing values. Panel **b** shows the ROC curves for the prediction performance on imputed data (black line) where we predict on the TCGA data with 5 genes randomly removed and the complete TCGA data (red line).

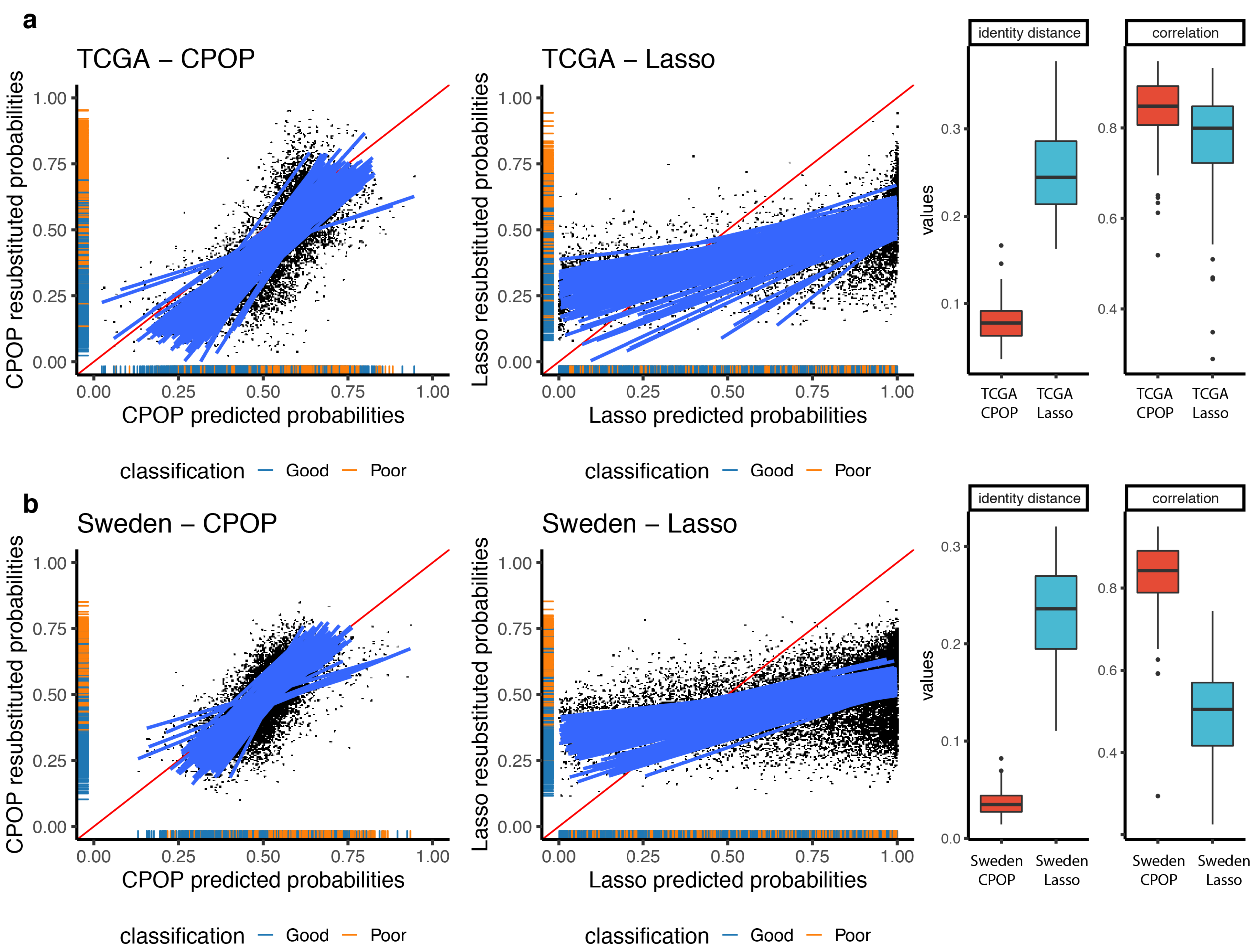


Supplementary Figure 12: We perform 20 repeated 5-fold cross-validation to select 100 different sets of features similar to Supplementary Fig. 5. We then plot concordance performance metrics from Supplementary Fig. 1 here. All feature sets are then re-fitted on the full datasets using ridge regression (similar to Fig. 2a). We compare the CPOP between-data predicted probabilities (x-axis) and the within-data re-substituted probabilities (y-axis) as prognosis probabilities for TCGA and Sweden data in panel a and b respectively. We also show the 100 linear regression models (blue lines) between the two. For a stable procedure like CPOP, we see that these regression lines are much more closely aligned with the identity line of $(y = x)$. As expected, CPOP offers better predictive performance than Lasso as well as having a good concordance with the re-substituted values.

### Supplementary Tables

- Supplementary Table 1 (separate CSV file): Table of NanoString panel genes for stage III melanoma prognosis.
- Supplementary Table 2 (below): Data collection and processing summaries for five melanoma datasets, MIA-Microarray, MIA-NanoString, TCGA, Sweden and MIA–Validation. Included are the data platform, number of samples, definition of the prognostic groups, data accession number, date of download and references. OS, RFS and DSS refers to overall survival, recurrence-free survival and disease specific survival, respectively.
- Supplementary Table 3 (below): Table of samples in the ovarian data, with median survival time.
- Supplementary Table 4 (separate CSV file): Table of CPOP coefficients for stage III melanoma prognosis constructed from MIA-Microarray and MIA-NanoString.

Supplementary Table 2: Data processing on five melanoma data.

| | Platform | n (good) | n (poor) | Definition of prognosis groups. | Processing strategies | Accession | Date | Reference |
|---|---|---|---|---|---|---|---|---|
| MIA - Microarray | Illumina Human WG-6 BeadChip microarray Version 3. | 19 | 26 | Good: RFS more than 4 years and alive with no sign of relapse. Poor: RFS less than 1 year and died due to melanoma. | NEQC normalisation | GSE54467 | 8 Jun 2016 | Mann et al. (2013); Jayawardana et al. (2015) |
| MIA - NanoString | Customised NanoString nCounter assay | 19 | 26 | Good: RFS more than 4 years and alive with no sign of relapse. Poor: RFS less than 1 year and died due to melanoma. | log2(raw counts) | GSE156030 | Between 24 May 2017 and 17 Apr 2018 | This paper |
| TCGA - SKCM | RNA-Seq | 43 | 34 | Good: survived more than the median survival time (26.9 mo). Poor: survived less than the median survival time. | log2(FPKM) | GDC-TCGA portal | 21 Apr 2018 | Network (2015) |
| Sweden | Illumina HumanHT-12 V4.0 beadchip | 67 | 64 | Good: survived more than the median survival time (17.6 mo). Poor: survived less than the median survival time. | Normalised values from GEO | GSE65904 | 12 Jan 2020 | Cirenajwis et al. (2015) |
| MIA - validation | Customised NanoString nCounter assay | - | - | - | | GSE156030 | Between 5 Mar 2020 and 13 Mar 2020 | This paper |

Supplementary Table 3: Samples and median survival time for nine ovarian data.

| | Data source | Abbreviation | Median survival (months) | Number of samples |
|---|---|---|---|---|
| 1 | Yoshihara et al. (2012) | Japan A | 104 | 260 |
| 2 | Tothill et al. (2008) | Tothill | 72 | 276 |
| 3 | Yoshihara et al. (2012) | Japan B | 96 | 40 |
| 4 | Bell et al. (2011) | TCGA RnaSeq | 73 | 557 |
| 5 | Bonome et al. (2008) | Bonome | 97 | 185 |
| 6 | Mok et al. (2009) | Mok | 52 | 53 |
| 7 | Yoshihara et al. (2010) | Yoshihara2010 | 76 | 110 |
| 8 | Bentink et al. (2012) | Bentink | 76 | 129 |
| 9 | Crijns et al. (2009) | Crijins | 52 | 157 |

### Additional Material A: Additional remarks on CPOP

In some situations, $\mathcal{S}^{(1)}$ in step 2 of CPOP might not select enough predictive features as some versions of the Elastic Net models have a tendency to only select one of many correlated features and ignoring the rest (Zou and Hastie, 2005). The most notable example of this is the Lasso (Tibshirani, 1996). To overcome this, we can enlarge this feature set by introducing an iterative component. This can be done by first calculating $\mathcal{S}^{(1)}$ as described above and then removing these features from the log-ratio matrices to obtain $\boldsymbol{Z}_{1,\mathcal{S}^{(1)c}}$ and $\boldsymbol{Z}_{2,\mathcal{S}^{(1)c}}$, where $\mathcal{S}^{(1)c}$ is the set complement of $\mathcal{S}^{(1)}$. We can then fit WEN onto $(\boldsymbol{Z}_{1,\mathcal{S}^{(1)c}}, \boldsymbol{y}_1)$ and $(\boldsymbol{Z}_{2,\mathcal{S}^{(1)c}}, \boldsymbol{y}_2)$ and update the feature set by adding the selected features in each iteration into $\mathcal{S}^{(1)}$. The removal of selected features in future iterations means informative correlated features are more likely to enter the feature set. The size of $\mathcal{S}^{(1)}$ is non-decreasing and empirically we find that 20 iterations are usually enough for the size of $\mathcal{S}^{(1)}$ to stabilise. In this paper, we primarily focus on the use of mean-difference weights: $w_j = |\,\text{mean}(\boldsymbol{Z}_{1j}) - \text{mean}(\boldsymbol{Z}_{2j})|$ for each $j = 1, \ldots, q$, whereas other choices are also available in our `CPOP` package.

### Additional Material B: Challenges in normalisation between datasets - adjusting for difference in scale

Given two arbitrary omics data measuring the same set of features, there is a very small chance that these two data will have equal scale. Fig. 1c provides an illustration showing boxplots of five melanoma gene expression data with four clear differences in scale. This is because omics features are typically measured on a relative scale with unit-less numerical values that are proportional to some molecular units. Technical batch effect across omics data of different origins is a classical example of this inconsistency and its presence in data is a key reason as to why omics data of different origins cannot be readily combined for the implementation of a clinical prediction work. Failing to correct for this scale difference in the data can produce misleading interpretation in the final predicted values.

For example, in Supplementary Fig. 4, we compute a Lasso model from samples in MIA - Microarray, MIA - NanoString, TCGA stage III samples (Network, 2015) and Sweden (Cirenajwis et al., 2015) samples. Then, for the TCGA stage III samples and Sweden samples, we compute the re-substituted prediction values. These values, drawn on the y-axis, are considered as gold standard of what the estimated risk probabilities should be from independent data. The prediction values from MIA - Microarray-trained Lasso model and MIA - NanoString-trained Lasso model are then plotted on the x-axis and coloured. Clearly, depending which data is used in the training of the Lasso model, we may arrive at different scale in the prediction. This comparison is particularly illustrative of the challenge associated with prediction in the presence of gene scale difference, as Fig. 1c shows MIA - Microarray and MIA - NanoString share high concordance for matched samples and yet their prediction values can show a large amount of variation.

Existing statistical methods aim to resolve gene scale differences through normalisation which inevitably requires estimation of data specific scales (e.g. mean and median) across all patient samples and a way to combine the samples. This procedure is often restrictive, since in a prospective experiment, it is not clear how a single sample should be combined with an existing study cohort (e.g. samples from previous studies frozen as a reference bank). Combining a single-patient's data with different cohorts and performing normalisation could lead to different predicted values for the same patient depending on the choice of the cohort.

The most popular method for adjusting data scales between different datasets is through the use of normalisation and this type of method can be divided into two broad categories: $z$-score standardisation and between-data normalisation. We refer to $z$-score standardisation as a pre-processing step where each omics predictor is centred at 0 and its sample variance is scaled to 1. This procedure has been used in, for example, Kossenkov et al. (2019); Deng et al. (2018); Liu et al. (2017); Chen et al. (2008); Herold et al. (2018); Waldron et al. (2014). On the other hand, between-data normalisation aims to correct the statistical distributions of genes in validation data to be similar to that of training data. Rudy and Valafar (2011) provides an evaluation of nine different between-data normalisation methods and more recent updates on this kind of normalisation include Taroni and Greene (2017) and Thompson et al. (2016). Through the act of combining multiple data and calculating summary statistics like sample-wise mean, both of these methods introduce interdependencies between the samples and may not be suitable for processing and predicting on a single omics profile (McShane et al., 2013, Criterion 15). Though within-sample standardisation methods such as in Lê Cao et al. (2014) have the potential to bypass some of these constraints, in this manuscript, we will operate under the assumption that re-normalisation for incoming samples together with existing cohort is not practical due to constraint with consent, data security or other reasons.

Data normalisation alone is not enough to address the various challenges in implementing precision medicine utilising omics data with additional considerations relating to the model stability and reproducibility also being necessary. While there are numerous modelling approaches we could have taken, for example, rank-based approach such as Eddy et al. (2010) and Afsari et al. (2015) or tree-based approach like Breiman (2001) and Ishwaran et al. (2008); we ultimately opt for a regression-based modelling approach because we want to efficiently utilise all gene expression information rather than summarising into ranks and wish to place a greater emphasis on model interpretability.
